# Cell therapy as a new approach on hepatic fibrosis of murine model of *Schistosoma mansoni* infection

**DOI:** 10.1101/853143

**Authors:** N. ALsulami Muslimah

**Affiliations:** Biology Department, Faculty of Science, University of Jeddah, Jeddah, Saudi Arabia

**Keywords:** BMSCs, histopathology, *Schistosoma mansoni*, OV-6

## Abstract

Schistosomiasis is an acute and chronic disease caused by blood flukes (trematode worms) of the genus *Schistosoma*. Schistosomiasis is disease that are prevalent in or unique to tropical and subtropical regions. Previous studies have shown that the role of bone marrow mesenchymal stem cells (BMSCs) therapy in improvement of hepatic fibrosis. Therefore, the current study was designed to assess the therapeutic role of BMSCs in murine schistosomiasis mansoni. BMSCs derived male mice were intraperitoneal injected into female mice that received *S. mansoni* cercariae through subcutaneous route. Mice were divided into four groups: negative control group (noninfected non treated); positive control group (infected non treated); BMSCs treated group; and untreated group. Liver histopathology and immunohistochemically were evaluated. BMSC intraperitoneal injection resulted in a significant reduction in liver collagen, granuloma size, and significant increase in OV-6 expression in the Schistosomiasis treated mice group. There was overall improvement of the pathological changes of the liver. The findings support that BMSCs has a regenerative potential in the histopathology and function of the liver tissue by decreasing liver fibrosis.

## Introduction

Schistosomiasis is caused by a trematode of Schistosomiasis and it is considered as a chronic debilitating disease. Schistosomiasis is considered the second disease after malaria infection in terms of its importance to public health, economic, and social effect. Estimates show that at least 207 million people infected all over the world, while 779 million people are still at risk (**Santiago et al., 2014**).

World Health Organization (WHO) proved that more than 200,000 deaths due to schistosomiasis each year in sub-Saharan Africa (**WHO 2012**). The most common cause of schistosomiasis in human is infection by *Schistosoma mansoni*. The disease is caused due to formation of a granulomatous reaction around the eggs of the parasite in the liver tissues (**Gryseels et al., 2006**).

Cirrhosis occurs as a result of chronic liver disease with various causes, including schistosomiasis which considered the most common cause of cirrhosis. One of the experimental models of liver fibrosis was infection with *S. mansoni*. It is used to illustrate the mechanism of the fibrosis (**El-Mahdi et al., 2014**).

Since 1970, the drug of choice is praziquantel (PZQ), which has proven effective in the prevention of some species of schistosomes, such as *S. mansoni*, *S. hematobium*, and *S. japonicum* and reduction of worm burden in infected patients. It is a safe oral drug that reduces the prevalence of schistosomiasis (**Eissa et al., 2011**). Therefore, the targeted and collective drug management program is currently heavily dependent on this drug to control morbidity induced by schistosomiasis (**Inobaya et al., 2014**).

With the choice of only one drug for treatment and the development of parasite resistance, the current situation is serious. Therefore, there is a real need to discover a new drug to mitigate the damages caused by liver fibrosis which induced by schistosomiasis (**Fikry et al., 2016**).

Previous studies reported that stem cells have a therapeutic role in tissue regeneration and require minor incursion procedures. They have used it lately because they have the capability of self-renewal and multilineage differentiation, which ably to diseases affecting human (**Salama et al., 2010**). It has been reported that transplanted bone marrow mesenchymal stem cells (BMSCs) lead to liver cell regeneration and perform a significant influence on liver structure and function (**Aziz et al., 2012**).

So, the destination of the current study was to assess the therapeutic role of BMSCs in the murine model of *schistosomiasis mansoni*.

## Materials and Methods

### Materials

- Reagents for cell culturing were purchased from Lonza Company, Swiss. They included:

▪ Dulbecco’s modified Eagles medium (DMEM) contained 4.5g/L glucose and L-glutamine (Catalog NO.: 12-604Q).
▪ Fetal bovine serum (FBS) (Catalog NO.: 77227).
▪ Phosphate buffer saline (PBS): without Ca^+2^ and Mg sterile filtered for cell culture (catalog NO.: BEBP17-516Q).
▪ Penicillin streptomycin mixture in 100 mLs bottles (Catalog NO.: 17-602E).
▪ Trypsin-EDTA (Catalog NO.: 17-161E).
- Wizard Genomic DNA purification kit (Promega, Madison, WI, USA) (Catalog NO.: **A1120**).
- SRY gene Primer sequences: (forward 5′ CATCGAAGGGTTAAAGTGCCA-3′, reverse 5′-ATAGTGTGTAG-GTTGTTGTCC-3′) published sequences (**Lee et al., (2010)** (UniGene Rn.107239).
- Immune-histochemical staining for OV-6 using monoclonal anti-mouse OV-6 anti-human antibody. It was purchased from Life Technologies Corporation and R&D Systems (Catalog NO.: FAB2020G).

### Experimental design

In the current study, forty female BALB/c mice (aged 4 weeks, weighing 30-40 g) were used and housed at Theodor Bilharz Research Institute (TBRI), Egypt. The experiment was proceeded in according to applicable guidelines in TBRI’s Animal Research Committee for testing animals. Animals were divided into four groups (10 mice/group) in this study.

**Group I:** negative control group; healthy uninfected mice injected intraperitoneal with phosphate buffer saline (PBS) and sacrificed after twelve weeks.

**Group II:** positive control group; mice subcutaneously infected with 90 cercariae of *S. mansoni* and sacrificed after eight weeks. Cercariae were gained from infected *Biomphalaria alexandrina* snails at the TBRI.

**Group III:** mice were infected as group II and on the 8th week they were infused via intraperitoneal injection by BMSCs (2×10^6^ cells/mouse) which derived from male mice then sacrificed after four weeks (**Aziz et al., 2012**).

**Group IV:** mice were infected as group II and left without treatment then sacrificed after twelve weeks. The infection was confirmed by examination of stool for the presence of *S. mansoni* eggs after 7weeks from infection.

### Isolation and culture of BMSCs

In the current study, 6-weeks male BALB/c mice was used as a donor of BMSCs. The tibiae and femurs were flushed using complete media which formed of DMEM supplemented with 10% FBS. Bone marrow cells resuspended for 2-3 days in complete media and incubated at 37°C in 5% humidified CO_2_. When colonies become confluence, cultures were washed with PBS. Then the cultures were trypsinized with trypsin EDTA for five minutes at 37°C. Then centrifuged the suspension, viable and nonviable cells were counted using hemocytometer. The resulting cultures were considered the first-passage cultures (**Abdel-Aziz et al., 2011**). After the third passage the BMSCs were used for intraperitoneal injection in treated group.

### Determination of male derived BMSCs in the treated group by polymerase chain reaction (PCR)

The presence or absence of the sex determination region on the Y chromosome male (sry) gene in recipient female mice was assessed by PCR. It was determined by Genomic DNA prepared from the tissue homogenate of liver tissue from mice in each group. The PCR were as follows: incubation at 94°C for 4 min; 35 cycles at 94°C for 50 s, 60°C for 30 s for optimal annealing, and 72°C for 1 min for extension; 72°C for 10 min. PCR products were electrophoresed using 2% agarose gel electrophoresis and stained with ethidium bromide. Y chromosomes marker was expressed as trans-illuminated line of the amplified product appeared at 104 bp (**Hong et al., 2013**).

### Histopathology study

After proper fixation of liver specimens using 10% neutral buffered formalin the specimens were dehydrated, cleared in xylol, impregnated and then embedded in paraffin wax. Five microns sections were cut and mounted on glass slides. The consecutive slides were stained with the following stains:

Hematoxylin and eosin (H&E) for examination of histopathological changes in the liver (**Bancroft et al., 2013**). Mallory stain to evaluate collagen content of the liver specimens (**Bancroft et al., 2013**). Immune-histochemical staining for OV-6 to evaluate the effect of transplanted BMSCs in the liver tissue (**Hegab et al., 2018**).

### Scanning electron microscope examination (SEM)

SEM was used to analyze the ultrastructural of the adult male and female *S. mansoni* worms. After eight weeks, adult worms were recovered from both the hepatic portal system and the mesenteric veins. Adult male and female worms of *Schistosoma mansoni* samples that served as controls were washed from blood using PBS, fixed in a 10% glutaraldehyde and processed for examination using SEM (JEOL- JSM - 5500 LV) at the Regional Center of Mycology and Biotechnology, El Azhar University, Cairo, Egypt (**Eissa et al., (2011)**; **Kamel & Bayomy, (2017)**).

### Morphometric measurements

The following parameters were measured using the image analyzer computer system (Leica Imaging System Ltd., Cambridge, U.K.). For each parameter at random selection of ten fields/ section in ten sections for every mouse in each group at magnification x 400 in a standard frame of 24790 μm^2^. The following parameters were measured: Area percentage of collagen fibers, the circumference of granulomatous lesion, and the mean number of OV-6 positive brown cells.

### Statistical analysis

Analysis of data was performed using one way-ANOVA and post hoc using SPSS 21 computer Software. The results were expressed as Mean ± standard Deviation (SD). P < 0.05 was considered statistically significant.

## Results

### Scanning electron microscope examination

The adult male *S. mansoni* worm as visualized by SEM showed anterior region with the oral and ventral suckers. The body increases in width directly behind the ventral sucker and folds ventrally to form the gynaecophoric canal with numerous tubercles distributed along the body (Fig. 1A). The oval oral sucker showed three regions: a large anterior part and posterior part, both covered with sharp spines of variable size and at the bottom, the oral cavity (Fig. 1B). Ventral sucker bigger and more prominent than the oral sucker. Spines did not present in both surfaces (Fig. 1C). On the dorsal surface of male worm tubercles and tegumental ridges with several spines were seen (Figs. 1D, E). Adult female worms showed the oral, ventral suckers, the sensory papillae and the integrity of the tegument with no abnormality observed (Figs. 2A-D).

**Fig. 1:**
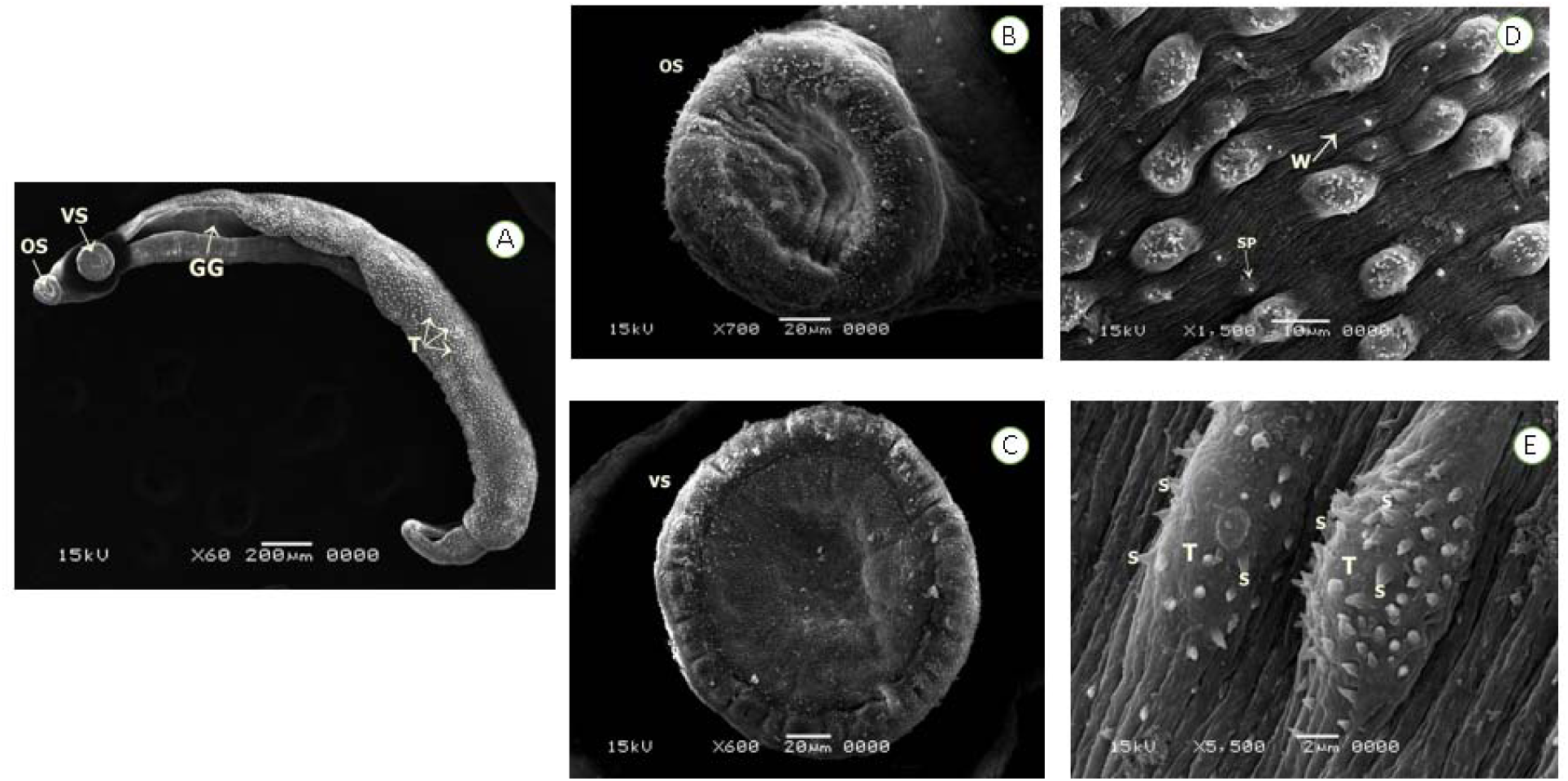
A: Male worms show the anterior portion of the body with oral (OS) and ventral suckers (VS) and the gynecophoral canal (GC) with no abnormalities. The dorsal region of male worms showing numerous tubercles (T) distributed along the body. B: superior and inferior borders of the oral sucker (OS). C: ventral sucker (VS) is bigger and more prominent than the OS. Both surfaces at this site did not present spines. D: in detail the dorsal region with sensory papillae (SP) and parallel wrinkles (W) visible. E: Higher magnification showing in detail numerous spines (S) covering the tubercules (T).

**Fig. 2:**
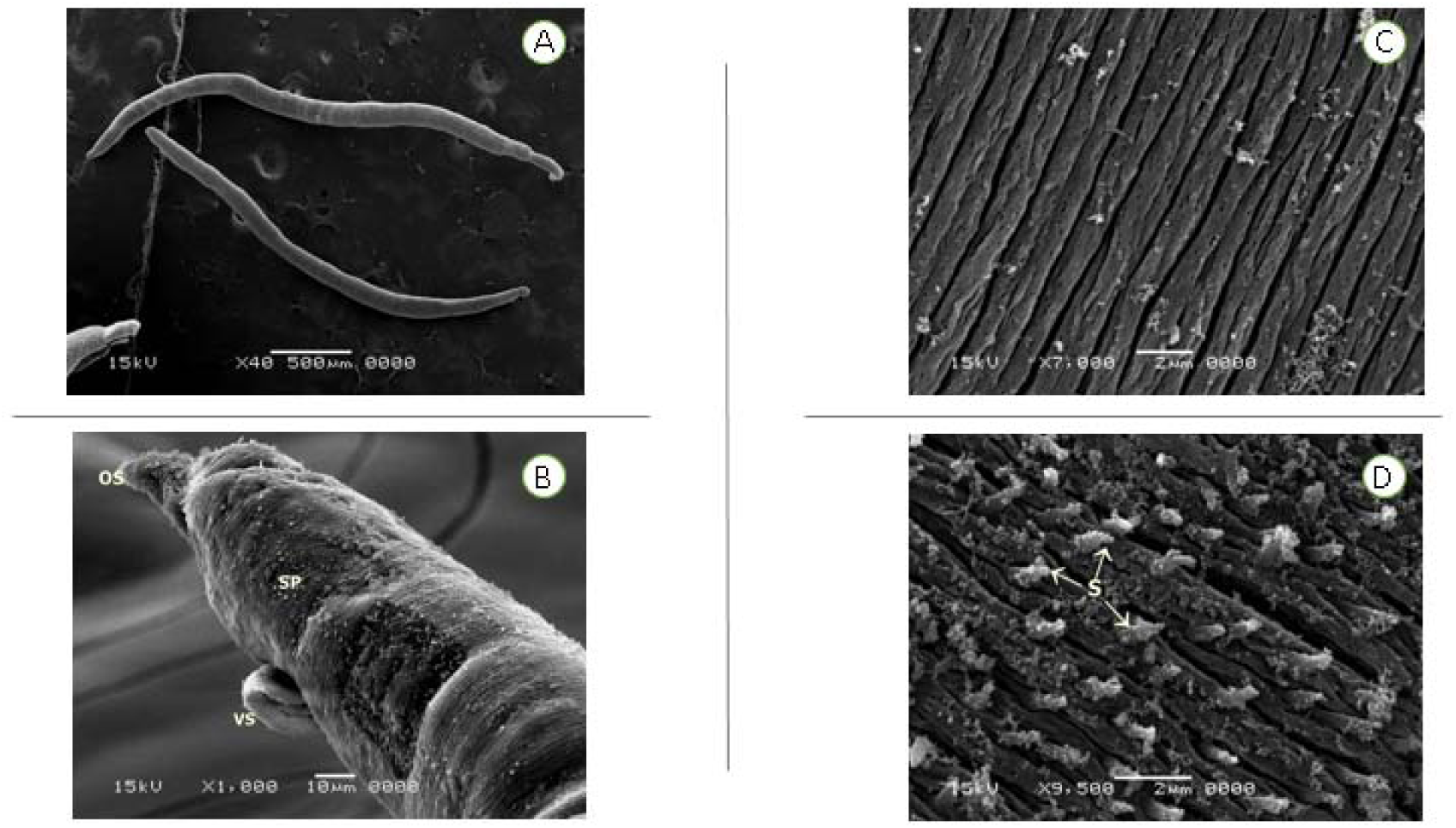
A: Female worm shows the whole extent of the body. B: Female worm showing the oral (OS), ventral (VS) suckers, the sensory papillae (SP) and the integrity of the tegument. C: spines (S) in detail. D: Higher magnification Female worms showing spines (S) in detail.

In the present study, the therapeutic potential of BMSCs on hepatic morbidity caused by *Schistosoma mansoni* was examined by culturing the undifferentiated BMSCs obtained from the femur and the tibia of male Balb/C mice with subsequent injection in the infected Balb/C female mice.

### Detection of male (Y chromosome) using PCR in the livers of female mice

Homing of injected BMSCs in the diseased liver was examined by PCR. PCR was done to detect the Sry gene (Y chromosome marker) in the mice groups which were injected with BMSCs. Therefore, the corresponding 104 bp band was detected only in test group III using Sry gene specific detection primers (Fig. 3).

**Fig. 3:**
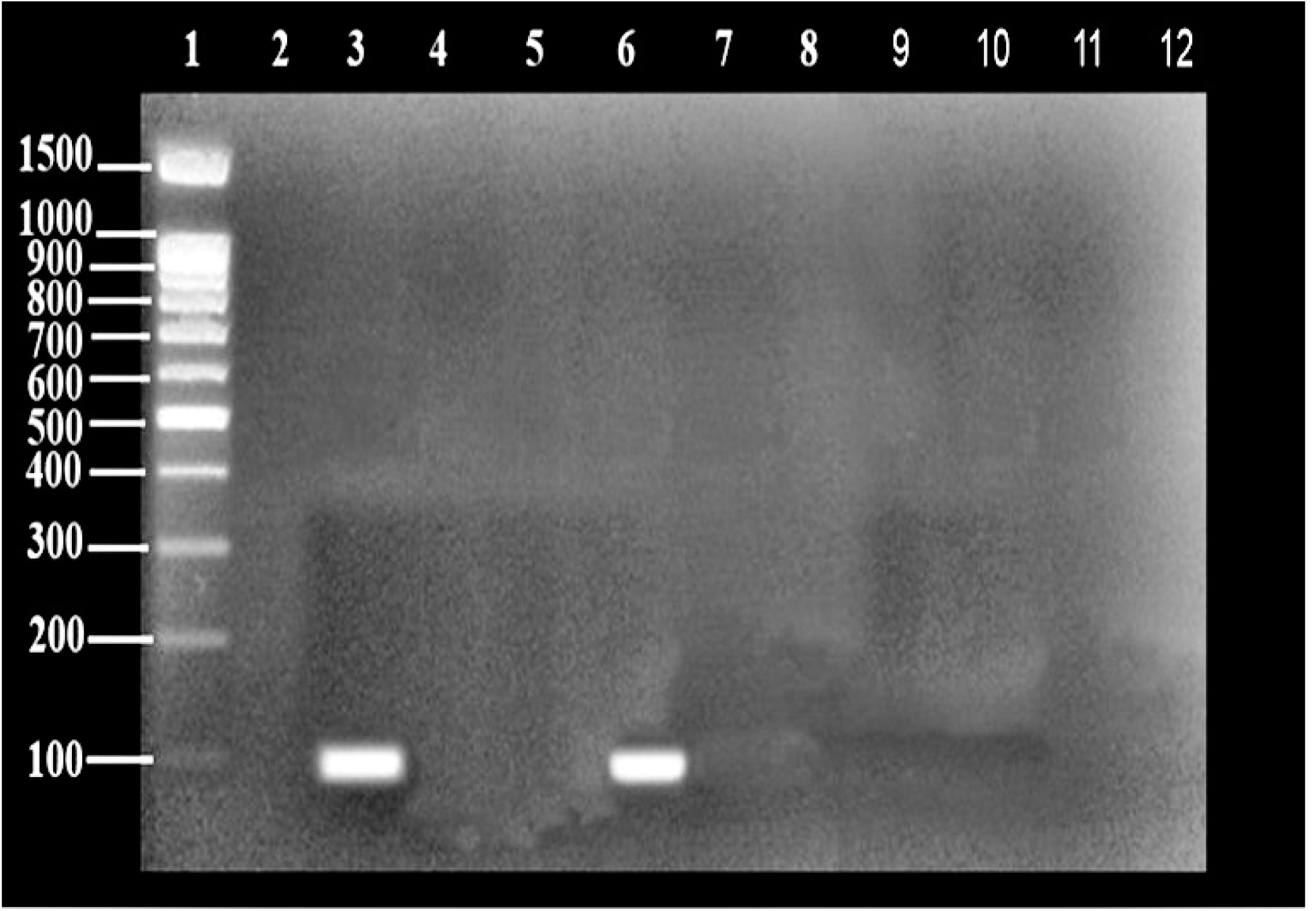
PCR analysis for Sry gene in liver homogenate of female mice groups**. Lane 1:** DNA molecular weight marker**. Lane 2:** Negative PCR control (female mice liver DNA). **Lane 3:** positive PCR control (male mice liver). **Lane 4:** Gr I. **Lane 5:** Gr II**. Lane 6:** Gr III. **Lane 7:** Gr IV.

Histopathological and morphometric, immunohistochemical and laboratory examinations were done to assess the effect of BMSCs injection on liver morbidity of the infected mice with *Schistosoma mansoni* (group III) to be compared with the group II and IV.

### Haematoxylin and eosin liver sections

The control group showed normal histological appearance of the classic hepatic lobule. Hepatocytes were noticed radiating from the central vein and separated by blood sinusoids. The hepatocytes appeared to be polyhedral acidophilic cells with a central rounded vesicular nucleus while few of them were binucleated. The portal tract at the corners of the hepatic lobule consisted of a branch of hepatic artery, portal vein and a bile duct (Fig. 4A-B). H & E stained sections of group III exhibited marked regenerative changes (Fig. 6A-D) compared to the infected non treated group II (Fig. 5A-C) and group IV (untreated group) (Fig7 A-D). These changes were demonstrated by reducing the infiltration of inflammatory cells, resolved pseudo-lobules and marked appearance of newly formed hepatocytes at the vicinity of the granuloma with a rounded dark nucleus.

**Fig. 4:**
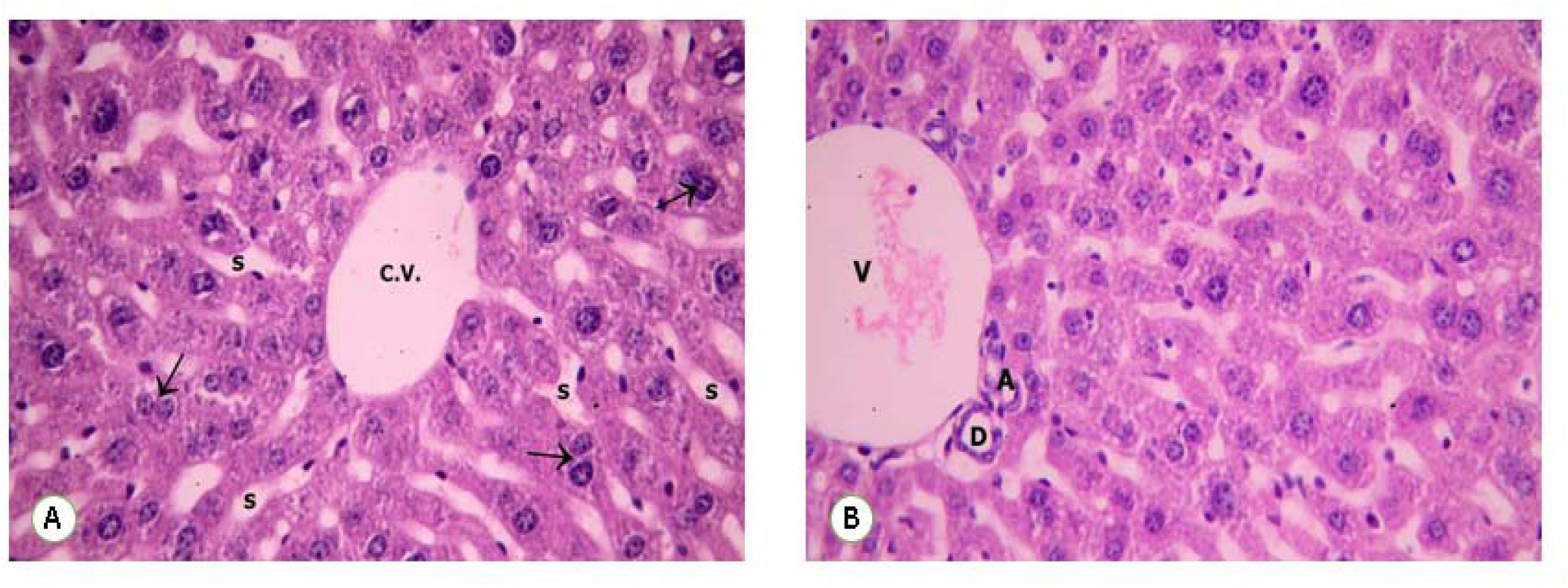
Photomicrographs of liver sections of control group I show: A: polyhedral hepatocytes with acidophilic cytoplasm and vesicular nuclei (↑). They are arranged as cords separated by blood sinusoids (S). These cords appear radiating from the central vein (CV). B: The portal tract containing a hepatic artery (A), portal vein (V) and a bile duct (D) can be also seen. H&E, x400

**Fig. 5:**
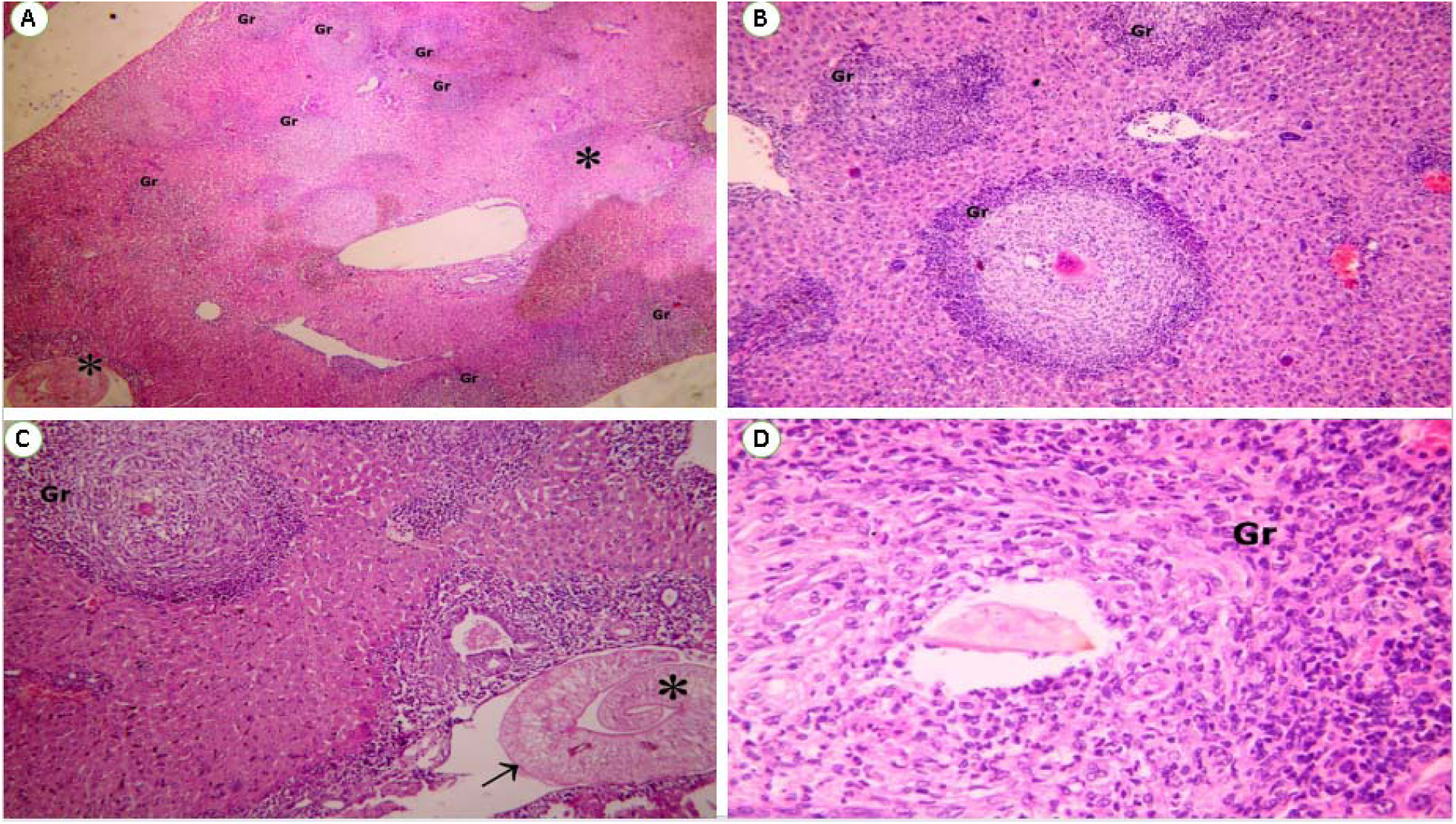
Photomicrographs of liver sections of group II (Schistosoma alone) show: A & B: multiple, large sized, cellular schistosomal granulomas (Gr) with inflammatory cellular infiltrate. H&E, x100, 200. C: Adults Schistosoma (*) shows the tuberculate exterior (↑) H&E, x200. C: the presence of large Schistosomal granuloma (Gr) around egg with marked inflammatory cells H&E, x400.

**Fig. 6:**
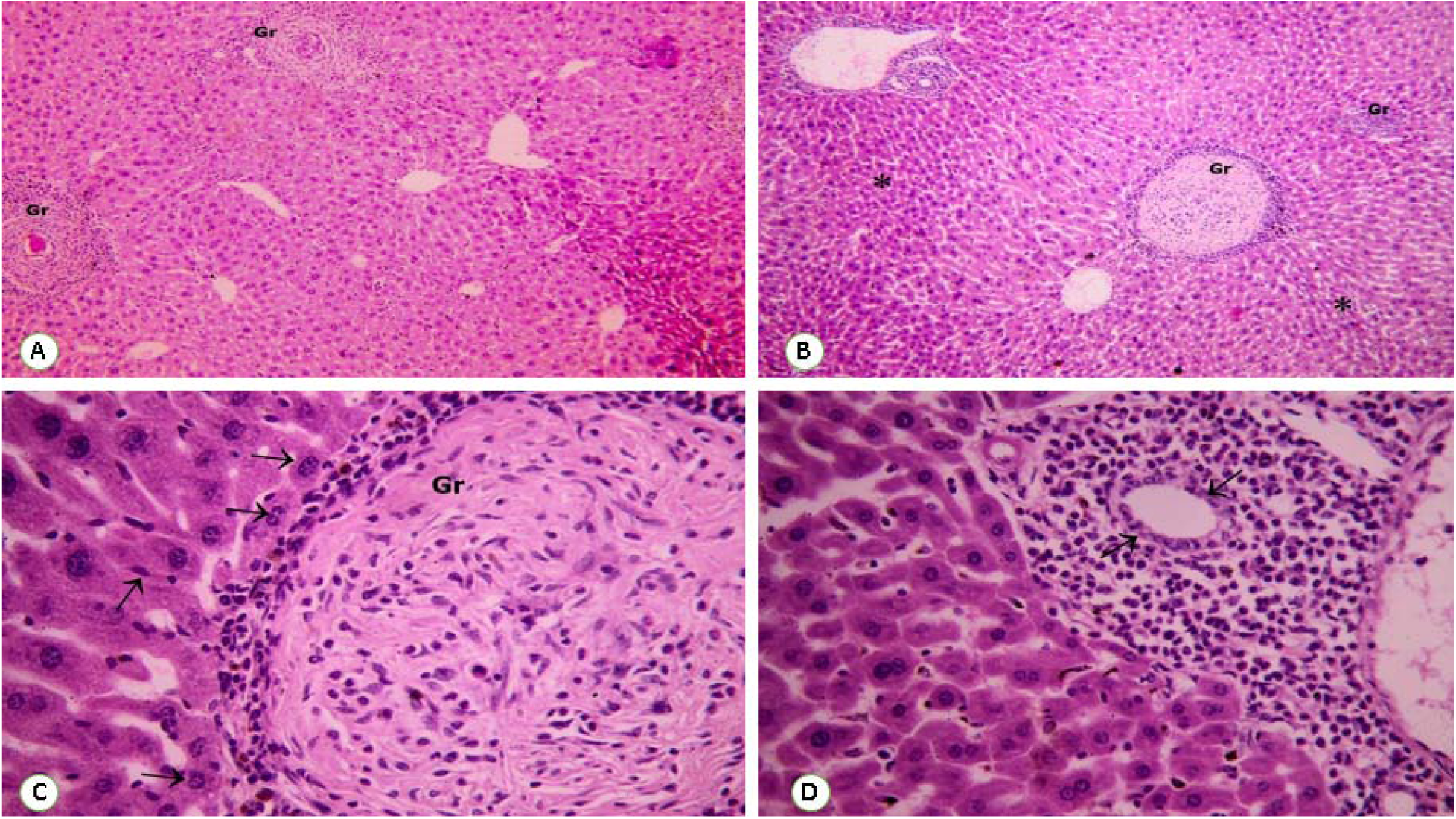
Photomicrographs of liver sections of group III (treated with BMSCs: A & B: show a significant reduction in granuloma size. Note the decrease of peripheral inflammatory cellular infiltrate with normal hepatic tissue in-between (*) H&E, x100, 200. C: show the presence of newly formed hepatocytic cells with relatively rounded to elongated large dark nuclei (↑) at the periphery of the granuloma (Gr) H&E, x400. D: show proliferating hepatic duct lined with columnar cells having basal nuclei (↑) H&E, x400.

**Fig. 7:**
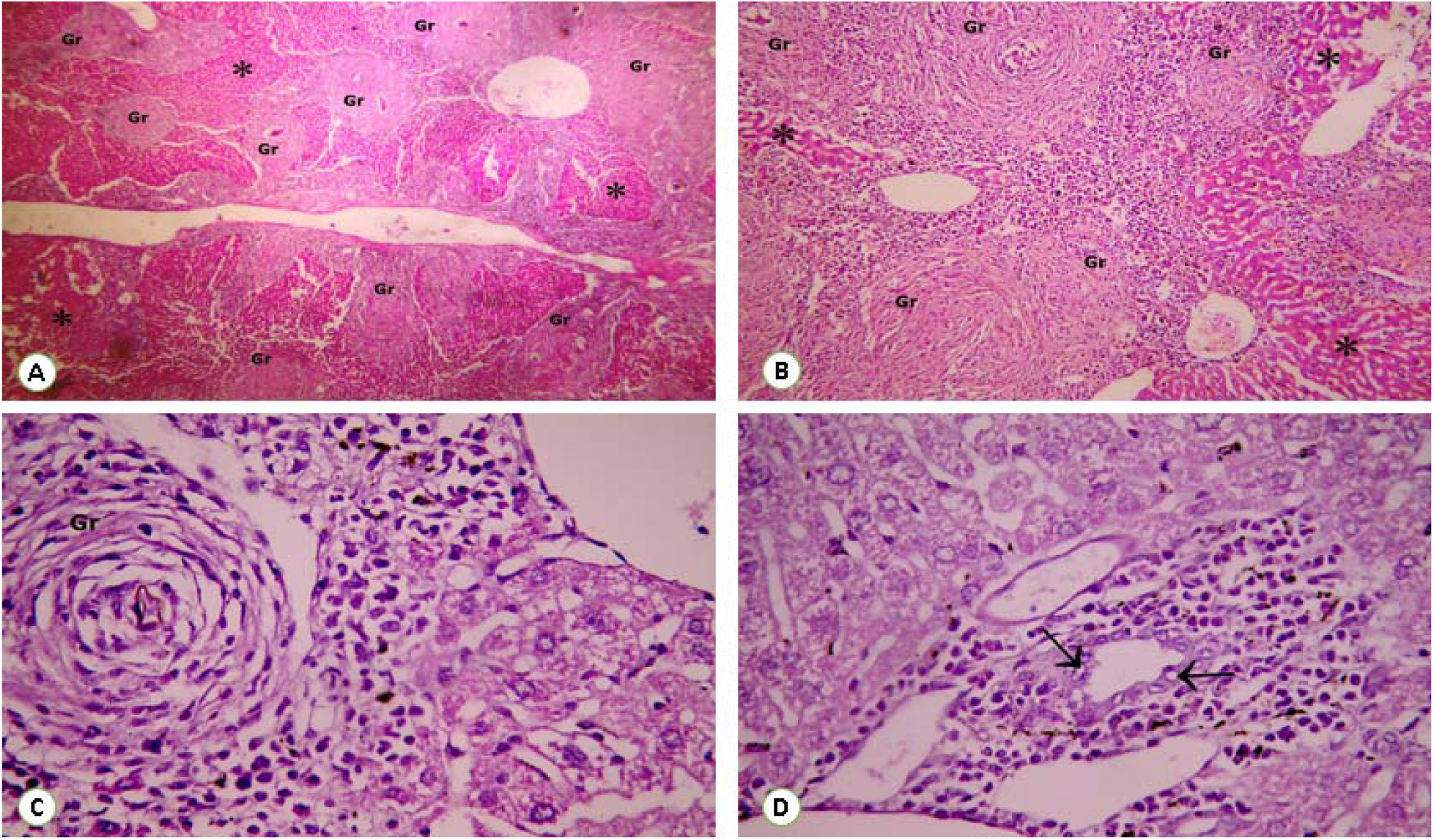
Photomicrographs of liver sections of group IV (Untreated group): A & B: show multiple coalescent characteristic fibrous schistosomal granulomas (Gr) with only few areas of hepatocytes (*) H&E, x100, 200. C: show absent of newly formed hepatocytic cells with calcified ova in the granuloma (Gr) H&E, x400. D: Show bile duct with highly vacuolated epithelium (↑) H&E, x400.

Morphometric analysis of granuloma sizes of H&E liver sections of Schistosoma-infected female mice of studied groups was assessed as shown in Table 1& fig. 8. There was a significant difference in mean area of the circumference of the granuloma size of group II, III and IV. There was a significant decrease in granuloma size of animal group III (treated with BMSCs) as compared to the group II (Schistosoma alone) and those mice of other groups IV (Untreated group). There was a significant increase was seen in animals of groups IV compared to groups II and III.

**Fig. 8:**
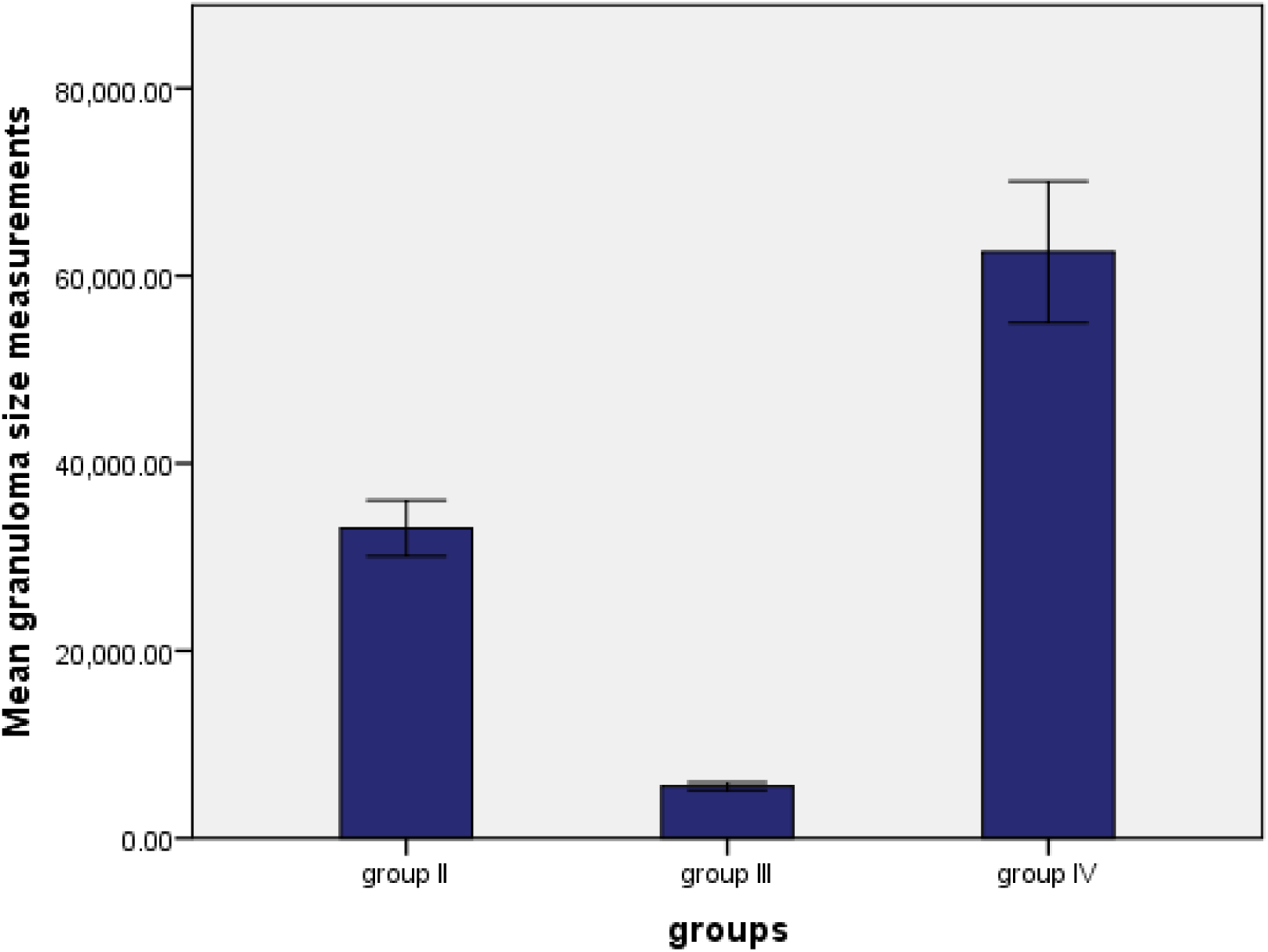
Comparison between the different groups as regard dimension of granuloma.

**Table 1:**
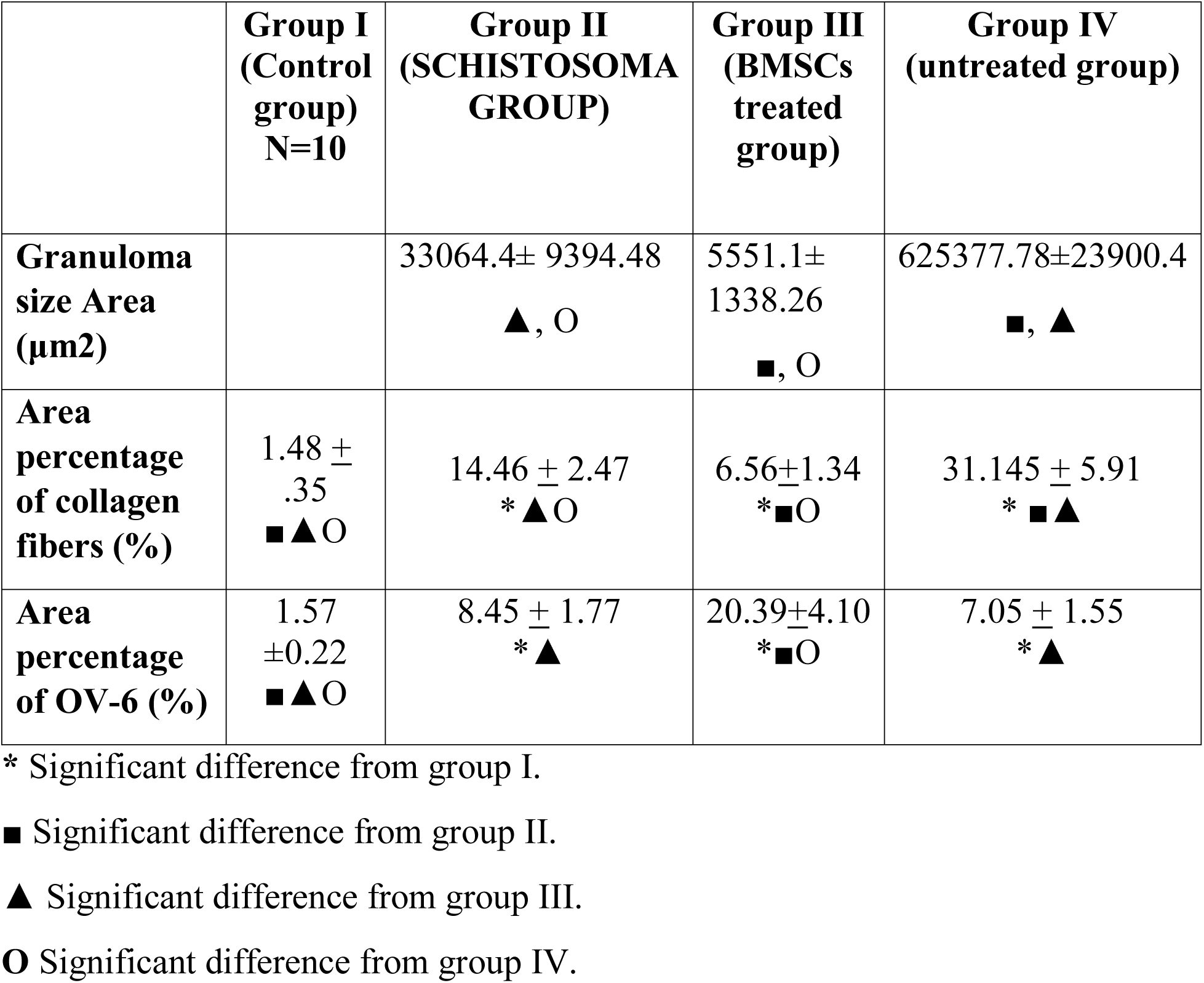
Comparison between granuloma size measurements (H&E sections), area percentage of collagen fibres (Mallory stained sections) and area percentage of positive OV-6 cells (IHC stained sections) in groups II, III and group IV

|  | **Group I**<br>**(Control**<br>**group)**<br>**N=10** | **Group II**<br>**(SCHISTOSOMA**<br>**GROUP)** | **Group III**<br>**(BMSCs**<br>**treated**<br>**group)** | **Group IV**<br>**(untreated group)** |
| --- | --- | --- | --- | --- |
| **Granuloma**<br>**size Area**<br>**($\mu\text{m}^2$)** | | 33064.4± 9394.48<br>**▲, O** | 5551.1±<br>1338.26<br>**■, O** | 625377.78±23900.4<br>**■, ▲** |
| **Area**<br>**percentage**<br>**of collagen**<br>**fibers (%)** | 1.48 ±<br>.35<br>**■▲O** | 14.46 ± 2.47<br>**\*▲O** | 6.56±1.34<br>**\*■O** | 31.145 ± 5.91<br>**\* ■▲** |
| **Area**<br>**percentage**<br>**of OV-6 (%)** | 1.57<br>±0.22<br>**■▲O** | 8.45 ± 1.77<br>**\*▲** | 20.39±4.10<br>**\*■O** | 7.05 ± 1.55<br>**\*▲** |

\* Significant difference from group I.
■ Significant difference from group II.
▲ Significant difference from group III.
O Significant difference from group IV.

### Mallory stain for liver fibrosis

Examination of Mallory stained slides showed few collagen fibers around central veins. Group II (Schistosoma group) showed multiple large granuloma with apparent increase of fibrosis around portal vein with impacted adult worm inside it. Group III showed significant reduction in the size of the granuloma and in area percentage of fibrosis. Group IV (Untreated group) showed multiple large schistosomal granuloma with apparent increase of fibrosis around portal vein and sparing only few areas of hepatic tissue (Fig 9 A-E). Group II (Schistosoma group) showed large schistosomal granuloma with marked inflammatory cells around Schistosoma egg. Group III showed granuloma with well-organized fibrous tissue, calcified Schistosoma egg and sparing most of the hepatic tissue. Group IV (Untreated group) showed large schistosomal granuloma with increased degree of fibrosis around Schistosoma egg (Fig 10 A-C). The histological results confirmed by statistical results of the area percentage of collagen fibers staining by Mallory stain. There was significant reduction in both group II and group IV compared to the negative control animals (group I). There was also significant decrease in area percentage of collagen fibers in group III compared to those mice of groups II and IV (Table 1 and Fig. 11).

**Fig. 9:**
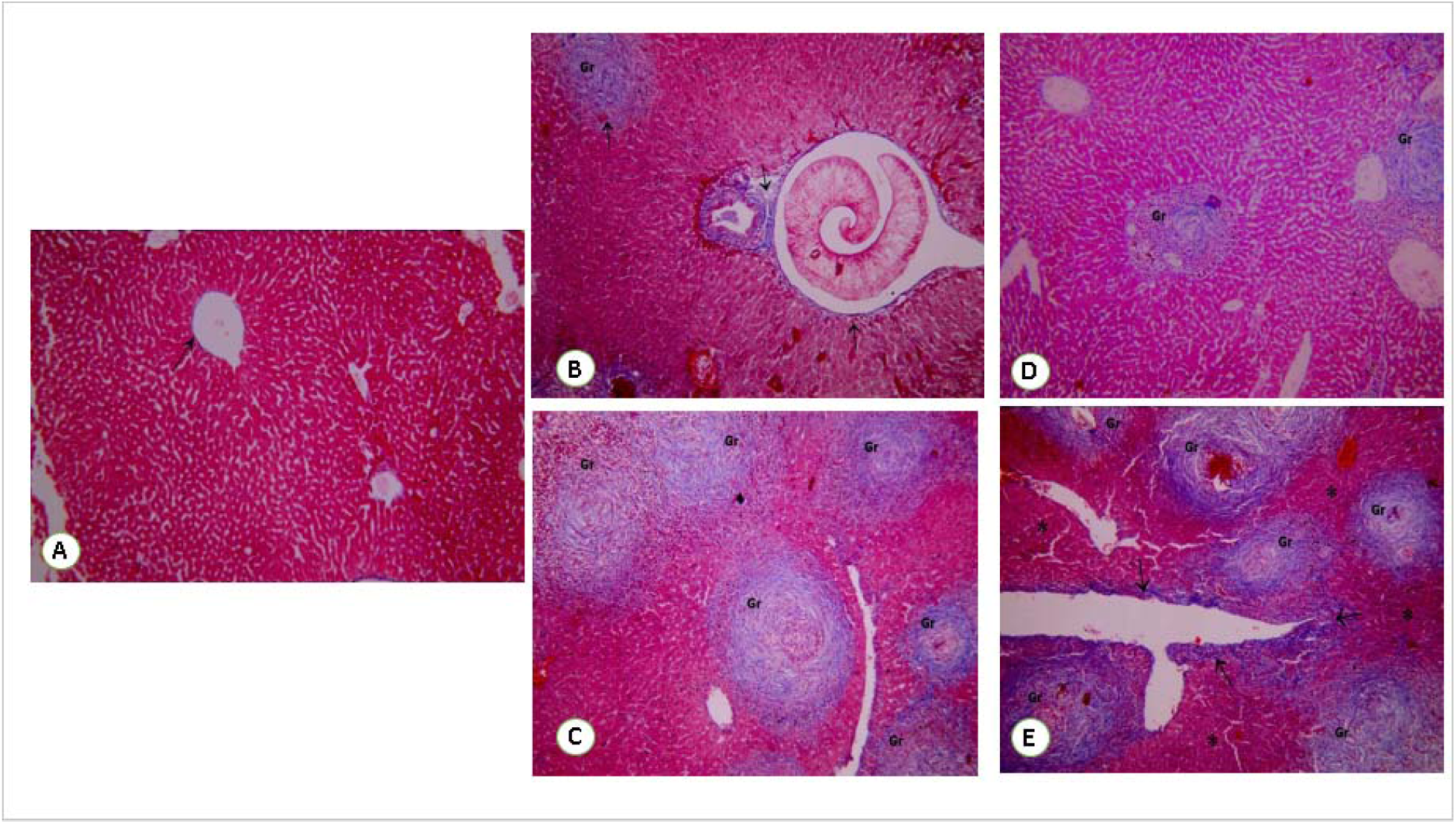
Liver tissue section from: A: Group I (control group), show few collagen fibers around central veins (↑). B & C: Group II (*Schistosoma* group) show multiple large coalescent schistosomal granuloma (Gr) with increased degree of fibrosis around portal vein with impacted adult worm inside it (↑). D: Group III (treated with BMSCs) show a significant decrease in granuloma (Gr) size and the degree of fibrosis. E: Group IV (Untreated group) show multiple large coalescent schistosomal granuloma (G) with increased degree of fibrosis (↑) around portal vein and sparing only few areas of hepatic tissue (*) **Mallory stain x100)**.

**Fig. 11:**
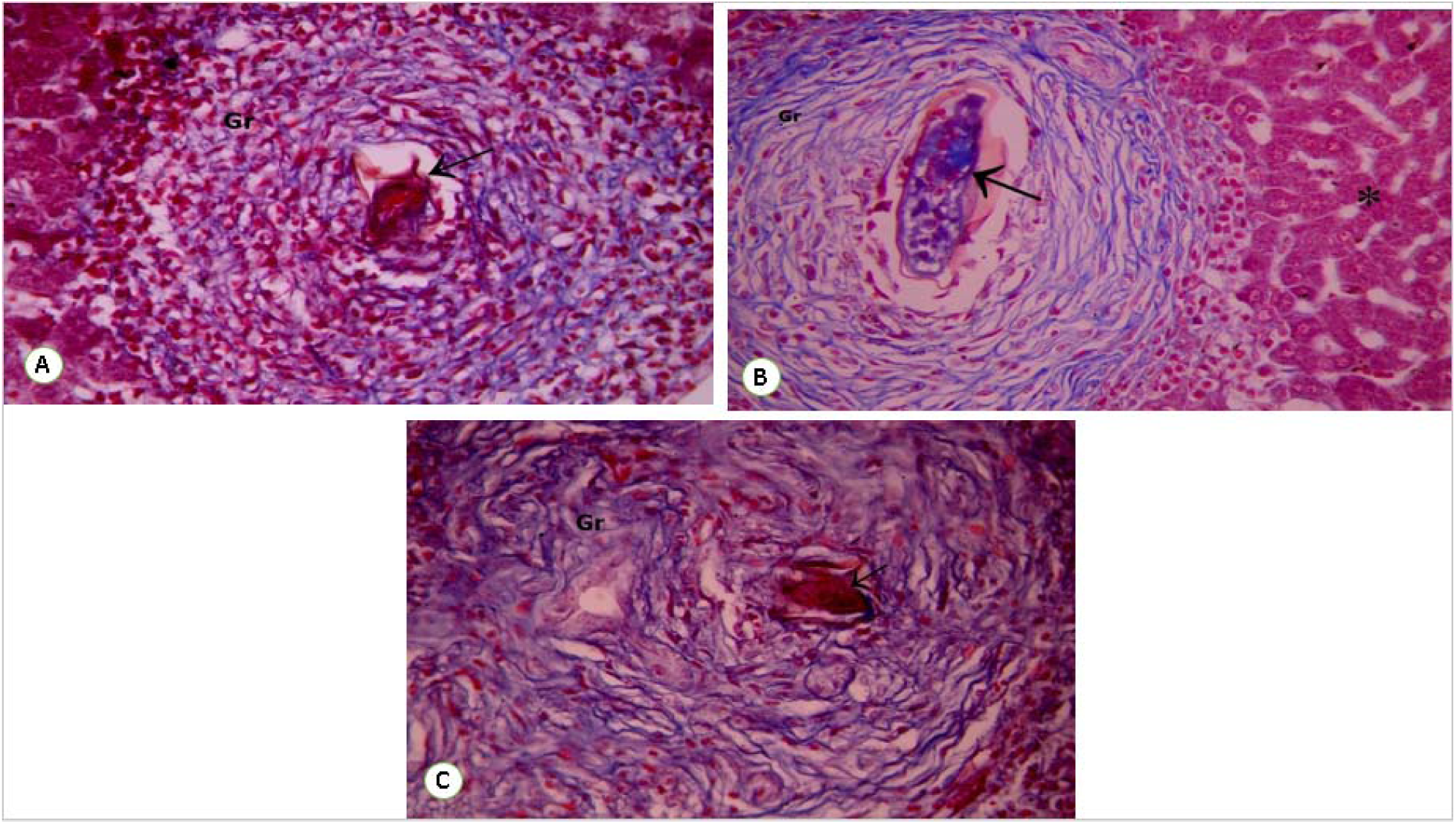
Liver tissue section from: A: Group II (*Schistosoma* group) show large schistosomal granuloma (Gr) with marked inflammatory cells and Schistosoma egg (↑). B: Group III (treated with BMSCs) show granuloma (Gr) with well-organized fibrous tissue, calcified Schistosoma egg (↑) and sparing most of the hepatic tissue (*). C: Group IV (Untreated group) show large schistosomal granuloma (Gr) with increased degree of fibrosis around Schistosoma egg (↑) **Mallory stain x400)**.

**Fig. 11:**
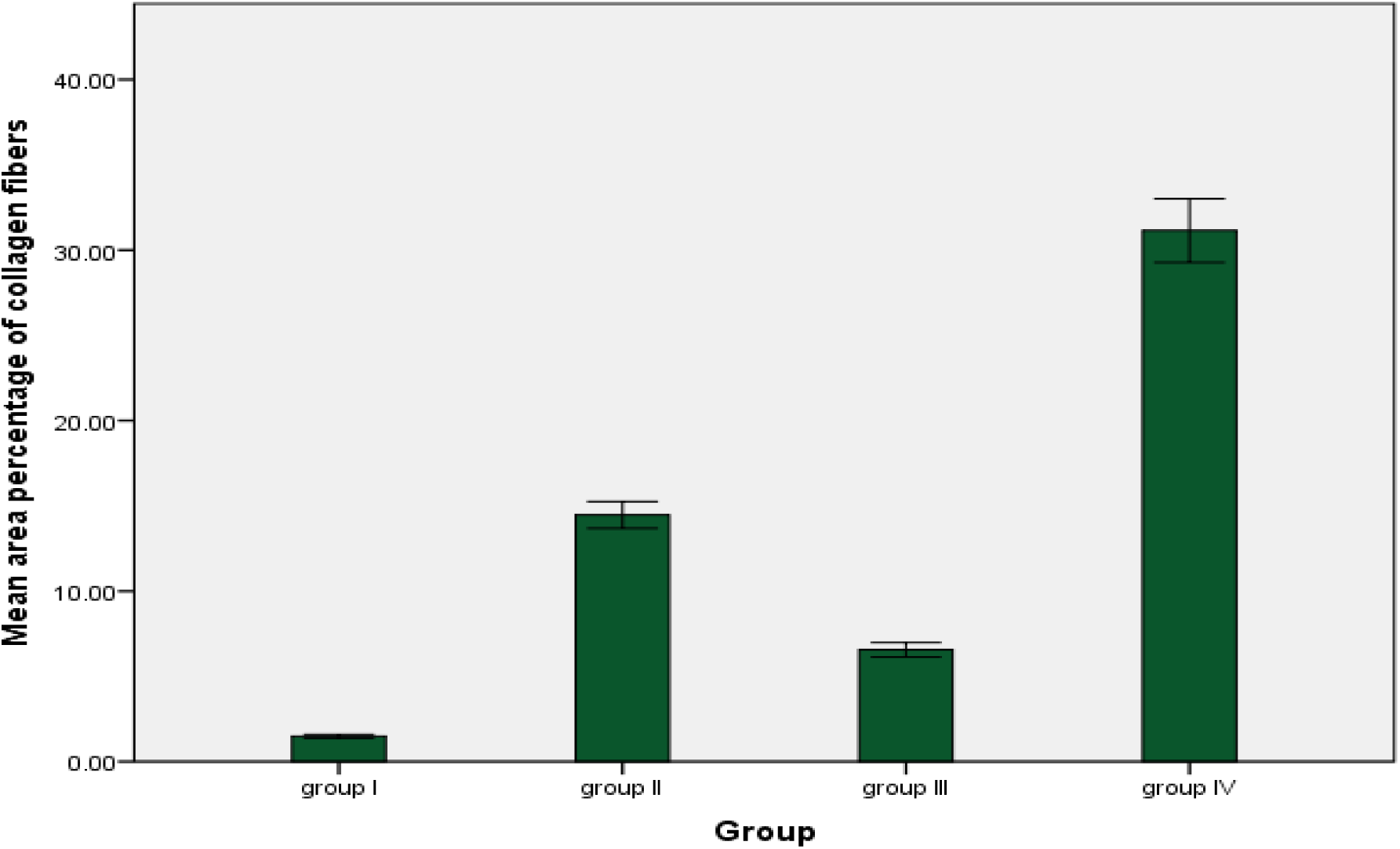
The mean values of the area percentage of collagen fibers among the different study groups.

### Immuno-histochemical detection of positive brownish OV-6 cells

Positive OV-6 cells in liver sections of negative control group, *S. mansoni* infected group (positive control group), group III (treated with BMSCs), and group IV (untreated group) were detected immunohistochemically. It is an indicator for the new formed hepatocytes differentiate from BMSCs. Negative control group exhibited no brown-colored positive OV-6 in liver cells. In the treated group with BMSCs, brown-colored liver cells appeared between mouse liver cells and around granulomas either individually or in groups. Group II (positive control group) and group IV (untreated group) showed few positive OV-6 in between liver cells and around the granuloma (Figs. 11A-E). The histopathological results of the current study were confirmed by statistical results. There was a significant difference in number of OV-6 positive cells in the different groups. They reached their highest peaks in treated mice with BMSCs (group III) as compared to the negative control mice and those mice of other groups II (positive control) and IV (untreated). Non-significant difference in OV-6 positive cells was seen in mice of groups II as compared to that seen in the group IV (Table 1 and Fig.13).

**Fig. 12:**
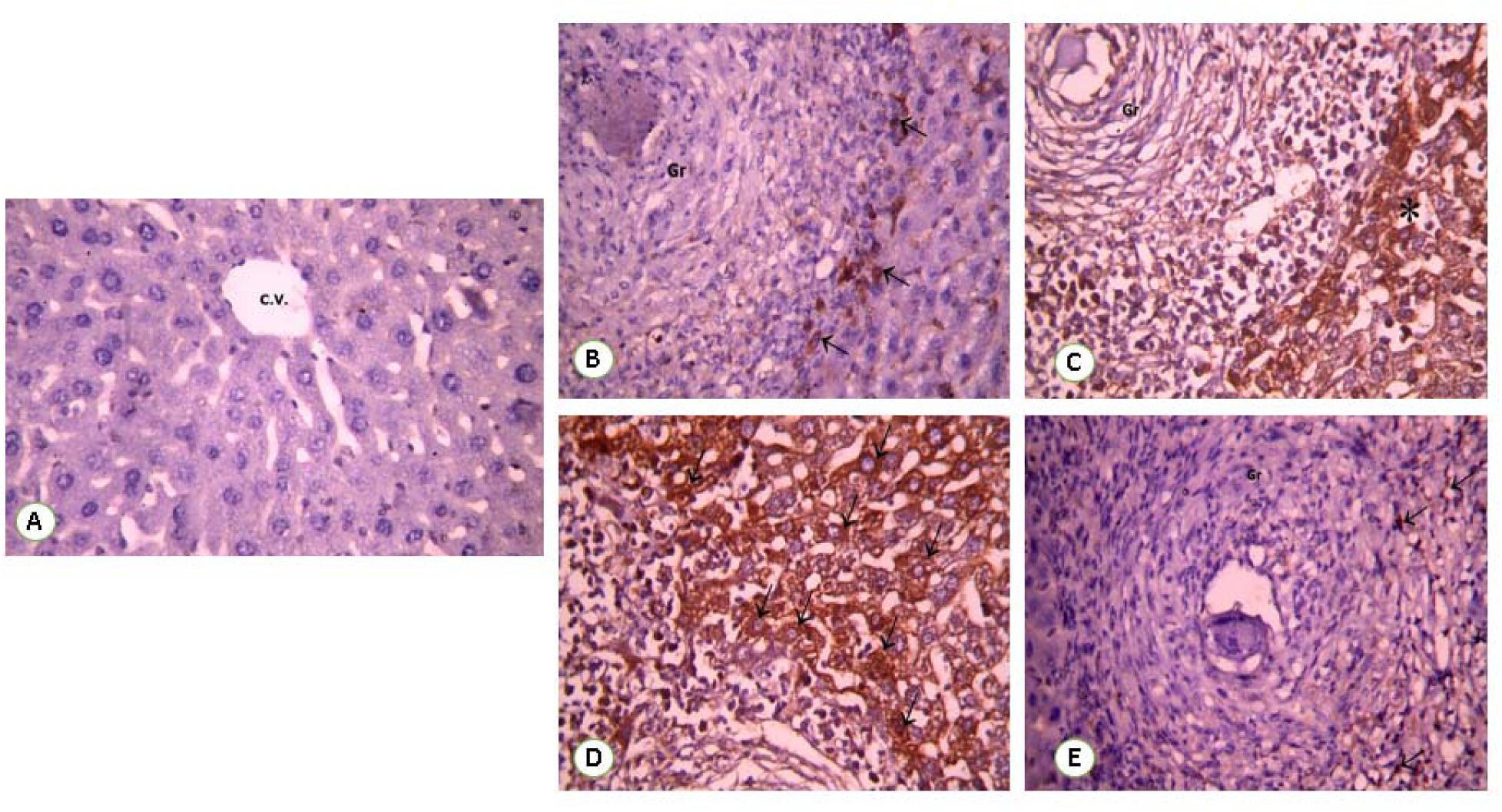
Immunostaining for OV-6 antibody in a liver section of: A: Group I (Control group) show negative expression of OV-6. Group B: show few positive expressions of OV-6 C &D: Group III (treated with BMSCs) show expression of OV6 monoclonal antibody as cytoplasmic brownish color (↑) in hepatocytes like cells (*) and granuloma cells (Gr). E: Group IV (Untreated group) show few positive expressions of OV-6 (↑) (IHC, DAB, **x400)**.

**Fig. 13:**
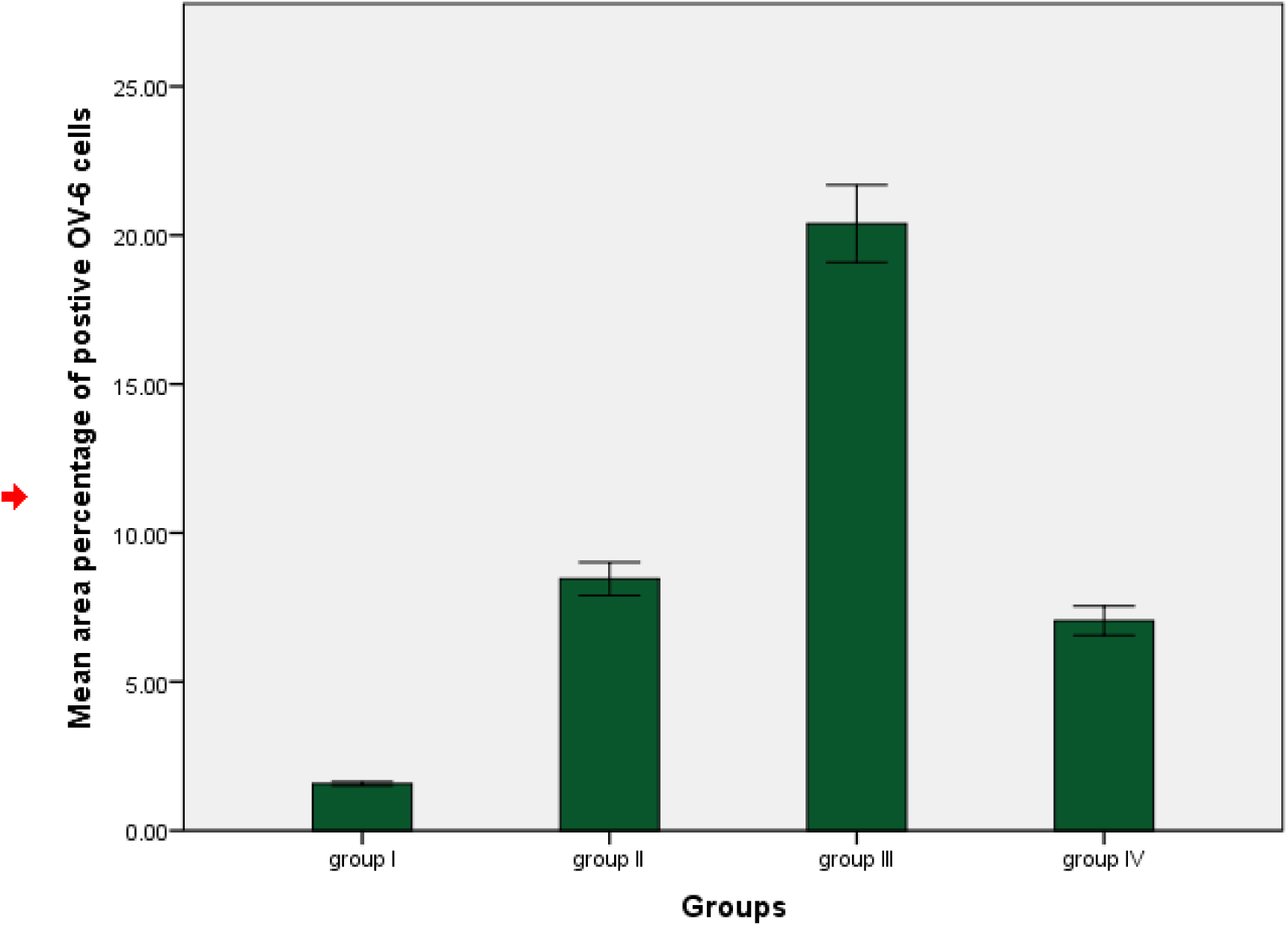
The mean values of the area percentage of positive cells of OV-6 among the different study groups.

## Discussion

Schistosomiasis is a chronic disease that is widespread all over the world. Schistosomiasis is endemic in 76 countries distributed all over the world especially in Africa and Asia (**Mata-Santos et al., 2014**). SEM plays a significant role in clarifying the detailed of the tegument of *S. mansoni* (**El-Shabasy et al., (2015)**; Kamel et al., (2017)). SEM have been used by various researchers to explain how anti-schistosomal drugs act on the adult worms (**de Oliveira et al., (2012)**; **El-Shabasy et al., (2015)**). The tegument is main interface between the parasite and the host (**El-Shabasy et al. 2015**). The thick tegument acts as an important drug target because the surface membrane function and the integrity of the tegument are essential for survival of the worm and its proliferation. These structures play vital roles in the immune evasion and nutrient absorption in the host (**Bertao et al., 2012**). In the present study, the male worms appeared with numerous tubercles with typical spines, oral and ventral suckers and the gynecophoral canal. Female worms were observed with parallel fissures and tegument spines. These findings are similar to findings observed by **Bertao et al., (2012) and Shabasy et al., (2015)** in their in vivo study.

The richest source of stem cells is the bone marrow (BM). BM derived stem cells (BM-DSCs) include mesenchymal stem cells (MSCs) and haemopoietic stem cells (HSCs). Enhanced understanding of these cells may help to develop novel therapies for regeneration of the diseased liver (**Puglisi et al., (2011)**; **De Miguel et al., (2019)**). **Wu et al., (2007)** added that an important reason for using the BMSCs therapy is their ability to home to injured tissues after administration. Another reason for using BMSC is their ability to differentiate into different types of tissues. When **Kinnaird et al., (2004)** cultured BMSCs, they were able to differentiate into the vascular components of the vascular bed as they enhanced the proliferation of endothelial cells.

Different authors reported the *in-vitro* differentiate of MSCs to hepatocytes (**Lee et al., (2004)**, **Cho et al., (2009)**; **Feng et al., (2011)**) and *in-vivo* (**Aurich et al., (2009)**; **Hu & Li et al., (2015)**). **Elkhafif et al., (2010)** transplanted BMSCs derived from male mice via intrahepatic route into infected female mice and reported new hepatocyte formation with reduction of granuloma size and fibrosis. Consequently, bone marrow represents the first source of MSCs because they can be easily obtained after aspiration from the sternum or the iliac crest of the human volunteer donors after informed consent (**Tondreau et al., 2004**).

In the current study, liver infection was induced by subcutaneous infection of immuno-competent Balb/C mice with 90 cercariae of *S. mansoni*. This technique was also performed by other authors as **Gryseels et al. (2006)**; **Elkhafif et al. (2010)**; **Hammam et al., (2016)**, who reported that inflammatory granulomas were formed inside livers of infected mice. **Gryseels et al. (2006)** described the acute liver granuloma on infected mouse (6-10weeks) post infection that appeared with different inflammatory cells, mainly eosinophils, lymphocytes, and macrophages. They also describe the chronic liver granuloma (16-20 weeks) post infection around the remains of the Schistosoma egg with dominant fibrous tissue.

In the present study, the regenerative capacity of the BMSCs was tested on murine model of S. mansoni infection. Intraperitoneal injection of BMSCs in *S. mansoni* infected mice lead to a significant reduction in the granuloma circumference and fibrotic areas with in-situ injection being the recommended route. This indicates that the inflammatory reaction and early fibrosis couldn’t hinder MSCs from reaching the injured liver (**Zhu et al., 2013**). Moreover, **Oliveria et al. (2008)** mentioned significant decrease of liver fibrosis after stem cell therapy that was occurred due to both intralobular and intravenous transplantation of stem cells. It was found that intravenous transplantation was more effective than other routs of administration such as intrasplenic transplantation, intraperitoneal, and intrahepatic routes (**Zhao et al., 2012**). In addition, **Xiang et al., (2005)** mentioned that the homing of BMSCs to the liver due to liver injury but not due to the site of injection of BMSCs.

In the current work, the injected BMSCs have been homed within the injured tissue irrespective of the route of administration. This was demonstrated by detecting Y chromosome of male BMSCs in liver tissue of group III by PCR. Previous researchers mentioned the appearance of injected BMSCs in injured liver (**Zhao et al., (2005)**; **Aziz et al., (2007)**; **Elkafif et al., (2010)**; **El-Mahdi et al., (2014)**; **Anan et al., (2016)**; **Fikry et al., (2016)**).

Different studies reported the regressive effect of BMSCs on liver fibrosis (**Zhao et al., 2012**). The mechanisms of the antifibrotic properties of BMSCs have been extensively studied; including inhibition of deposition of collagen fibers (**Mohamadnejad et al., 2007 and Horton et al., 2013**). Some authors reported increased levels of matrix metalloproteinase which may directly degrade the extracellular matrix and lead to apoptosis of hepatic stellate cell (HSCs) which is the cells responsible for fibrosis and collagen production in the liver tissue (**Zhao et al., 2005**). Recently, **Ahmed et al., (2014)** have reported that BMSCs were able to attenuate cirrhosis by a directly suppressing the activation of HSC, in an animal model of liver fibrosis induced by carbon-tetrachloride.

To confirm the regenerative findings in form of newly formed hepatocytes seen in H & E liver sections of treated BMSCs injected groups in the present work, immunohistochemical examination of OV-6 cells was done. It revealed the presence of newly formed hepatocytes in between the inflammatory cells at the vicinity of granuloma and also at the peripheral areas. They displayed a significant increase of positive cytoplasmic reaction of OV-6 cells. **Oliveira et al. (2008)** stated that increased 3-4 folds of oval cells were found in liver sections of mice treated with BMSCs which meant new hepatocytes formation. Similar findings were recorded by **Elkhafif et al., (2010)** who mentioned that the origin of the newly formed hepatocytes could be the injected BMSCs or from the liver itself (regenerated hepatocytes). **Alison et al., (2009)** reported that the improvements could be due to the secretion of growth factors by BMSCs, rather than to their differentiation into liver cells.

The present work showed high regenerative capacity of BMSCs as evidenced by the newly formed hepatocytes, the reduction of the inflammatory cells as well as the antifibrotic effect which were reflected on the granuloma size, the collagen fibers, OV-6 positive cells. This is also noticed by **Hegab et al., (2018)**. Moreover, **El-Mahdi et al., (2014)** stated BMSCs could differentiate into liver cells which lead to improvement of the remaining liver cells function after severe hepatic injury. Also, Oval cells could be differentiated into hepatocytes, cells of bile duct, and promote hepatic regeneration (**Wang et al., (2003)**; **Fausto and Campbell, (2003)**).

In conclusion, injection of BMSCs in S. mansoni-infected mice resulted in decrease granulomas’ size, regression of liver fibrosis and contributes to the generation of new hepatocytes. These findings reinforce the trial of BMSCs in patients with chronic and immunological liver diseases including schistosomiasis.

